# Plant–pollinator interactions along an elevational gradient in Central European forest understorey

**DOI:** 10.64898/2026.08.15.745048

**Authors:** Antigoni Sounapoglou, Štěpán Janeček, Sailee P. Sakhalkar, Ishmeal N. Kobe, Eliška Chmelová, Dominik Anýž, Sylvain Delabye, Jan Filip, Jiří Hodeček, Karolína Jackwerth, Renata Píplová, Kateřina Hanzelková, Tomáš Křížek, Yannick Klomberg, Jan E.J. Mertens, Robert Tropek

## Abstract

Elevational gradients provide a framework for understanding how environmental filtering reorganises communities and interactions, but plant–pollinator interactions along temperate forest elevational gradients remain overlooked. We studied early-spring understorey communities at four forest sites spanning the foothills towards the timberline (450–1,000 m a.s.l.) in the Krkonoše Mountains, Czechia. Across six transects per elevation, we quantified flowering plant species richness, floral resources and traits, and video-recorded flowers, yielding 4,003 pollinator visits. We analysed elevational patterns in species richness, community composition, floral traits, and quantitative network characteristics. Visitation frequency and flowering plant and pollinator species richness peaked at intermediate elevations. The contribution of dipteran relative to hymenopteran pollinators increased towards higher elevations, principally because of non-hoverfly flies, whereas individual bee groups showed no uniform response. Floral resources and traits showed no uniform elevational responses, although total nectar sugar availability peaked at the highest site because of the dominant *Vaccinium myrtillus*. Most notably, both network-level specialisation and mean species-level specialisation were generally greater at the two higher elevations, whereas nestedness was lower and other network characteristics showed no consistent patterns. These findings suggest that shifts in pollinator composition and dominant floral resources potentially shaped interactions along the gradient. The increasing specialisation with elevation contrasts with the generalisation often expected under reduced partner availability and indicates that forest networks may follow elevational patterns not predicted from open habitats. Despite limited site-level replication, this study provides, to our knowledge, the first community-wide characterisation of plant–pollinator interactions along a temperate forest elevational gradient and identifies patterns requiring evaluation across replicated gradients.

## Introduction

Elevational gradients shape ecological communities by filtering species according to their climatic tolerances, phenology, and resource requirements (Sundqvist et al. 2013). Species richness often decreases towards the upper limits of these gradients, although monotonic, hump-shaped, and weak or absent relationships have all been reported (Guo et al. 2013; McCain and Grytnes 2010). In the European Alps, plant species richness responded jointly to climate and management, with insect-pollinated plants reaching their species richness maximum under cooler conditions than wind-pollinated plants (Hoiss et al. 2013). For terrestrial insects, the main pollinator group, a recent synthesis identified a broad low-elevation plateau followed by declining species richness as a dominant elevational diversity pattern, although other patterns may occur as well (Dolson and Kharouba 2024). Importantly, plant and pollinator diversity need not respond synchronously because the two groups differ in their environmental tolerances, resource requirements, and phenology. As plant–pollinator interactions require the spatial and temporal co-occurrence of both partners, turnover within either group can alter the available interactions more than total species richness gradients alone (Minachilis et al. 2023).

Besides plant species richness, elevation can alter both the quantity of floral resources and their accessibility to different pollinators. Flower abundance is strongly affected by local productivity and phenology: in the French Alps, its relationship with elevation varied non-linearly through the season and flowering peaks occurred progressively later at higher elevations (Lefebvre et al. 2018), whereas surveys in the Austrian Alps found declining floral abundance with elevation (Aguirre and Junker 2024). Intraspecifically, however, a recent meta-analysis of 121 angiosperm species found no general elevational pattern in flower number (Novaes et al. 2025). Elevational patterns in floral morphology also appear strongly trait- and scale-dependent. Flower size showed no consistent elevational pattern in either the Swiss or Austrian Alps (Fabbro and Körner 2004; Junker and Larue-Kontić 2018), while floral tube length was likewise unrelated to elevation in the Austrian Alps, although tube width decreased with elevation (Junker and Larue-Kontić 2018). Intraspecific studies have likewise found both decreases and increases in floral dimensions with elevation, sometimes associated with changes in pollinator size and/or composition (Sun et al. 2014; Kuriya et al. 2015; Kiełtyk 2021), consistent with the absence of a general intraspecific global pattern in flower size in the recent meta-analysis (Novaes et al. 2025). Community-level information on nectar sugar production per flower or total nectar sugar availability in the community remains largely missing for temperate elevational gradients, to our knowledge, although these rewards directly define the resources for pollinators.

Pollinators are primarily filtered by the colder, wetter, and more seasonal conditions at higher elevations of temperate regions (Hodkinson 2005; Hoiss et al. 2012; Dolson and Kharouba 2024). One of the most widely reported elevational patterns is a shift from bee-dominated communities at lower elevations towards greater importance of flies under colder and wetter conditions (Kearns 1992; Lefebvre et al. 2018; McCabe et al. 2019; McCabe and Cobb 2021). Bees obtain both adult food and larval provisions from flowers, whereas flower-visiting flies have diverse larval diets that are generally unrelated to floral resources (Larson et al. 2001). Consequently, reduced flower availability may constrain bees more strongly than flies, while cooler environments favour many fly taxa (McCabe et al. 2019; McCabe and Cobb 2021). Nevertheless, individual groups can depart from this general pattern. In particular, bumblebees can remain important pollinators in cold mountain environments (Sponsler et al. 2022), and both wild bees and hoverflies remained abundant and species-rich at higher elevations in extensively managed hay meadows (Baumann et al. 2021). A greater contribution of flies at higher elevations is therefore more consistently expected than any uniform response across all bees.

Changes in plant resources and pollinator assemblages are jointly translated into the organisation of their interaction networks. The altitudinal niche-breadth hypothesis predicts broader realised interaction niches and lower specialisation at higher elevations, where greater environmental variability and altered resource availability may favour broader partner use (Rasmann et al. 2014). Conversely, if colder or more seasonal environments at higher elevations reduce temporal overlap among floral resources and shorten pollinator flight periods, specialisation may increase (Glaum et al. 2021). A recent global analysis likewise found that climate structured plant, pollinator and network-level specialisation in complex and taxon-specific ways rather than producing a universal geographic trend (Sakhalkar et al., in press). Connectance, network- and species-level specialisation, modularity and nestedness capture complementary aspects of how interactions are distributed and restricted among partners, but their elevational patterns may also partly reflect geometric constraints on species co-occurrence (Fibich et al. 2026). Temperate mountain studies have accordingly reported decreasing network-level specialisation in alpine grasslands, increasing nestedness and generalisation in high-elevation forest zones, and non-linear changes associated with the functional composition of plants and pollinators (Hoiss et al. 2015; Chesshire et al. 2021; Aguirre and Junker 2024).

Most evidence underlying these ideas in temperate regions comes from open habitats, such as meadows and other grasslands. Forests differ from open habitats in their canopy-buffered microclimate and in the amount and seasonal distribution of understorey floral resources, with many temperate deciduous-forest herbs flowering before canopy closure (Kudo et al. 2008; McCabe et al. 2019; De Frenne et al. 2019), limiting the transferability of results from open habitats. To our knowledge, the only temperate studies focused on pollinator communities or interaction networks along a forest elevational gradient come from the same mountain system in southwestern USA, and both cover only a limited montane section of its full elevational gradient (McCabe et al. 2019; Chesshire et al. 2021). We are unaware of comparable community-wide data spanning forest habitats from the lowlands towards the timberline in any temperate mountains. This deficiency reflects the wider neglect of forest pollinator ecology, including a pronounced shortage of detailed plant– pollinator interaction data from boreal forests (Ulyshen et al. 2023; Díaz-Calafat et al. 2025).

This study provides, to our knowledge, the first community-wide characterisation of plant– pollinator interactions along a temperate forest elevational gradient extending from the foothills towards the timberline. Focusing on the early-spring flowering peak in the understorey of the Krkonoše Mountains, Czechia, Central Europe, it combines continuous video recording of floral visits with measurements of floral traits and quantification of floral resources at the community level. We asked how elevation was associated with (i) plant and pollinator species richness, flower abundance, floral sugar availability, and visitation frequency; (ii) the representation of the principal pollinator functional groups; (iii) quantitative network structure and plant and pollinator specialisation; and (iv) distribution of selected floral traits, including flower size, floral tube length, and nectar sugar production. We expected the higher sites to support fewer plant and pollinator species and lower visitation frequencies than the lower sites, together with a greater relative contribution of flies compared to bees. We further expected both network- and species-level specialisation to decrease and connectance to increase with elevation, but had no directional expectations for modularity or nestedness. Given the inconsistent patterns reported previously, we made no directional predictions for floral traits, flower abundance, or floral sugar availability.

## Methods

### Sampling area and design

The fieldwork was performed on the southern slopes of the Krkonoše Mountains in northern Czechia. The mountains rise from the foothills at ∼400 m a.s.l., through the timberline at an average elevation of ∼1,240 m a.s.l., to the arcto-alpine tundra of their highest parts, reaching 1,603 m a.s.l. on Mt. Sněžka (Flousek et al. 2007; Treml and Migoń 2015). The region has a relatively cold and wet temperate climate, with mean annual temperatures declining from ∼6 °C at the foothills to ∼2 °C on the highest ridges, and annual precipitation increasing from ∼800 mm at the foothills to 1,200–1,400 mm along the highest ridges (Flousek et al. 2007). The Krkonoše Mountains southern slopes are covered predominantly by a cultural landscape comprising forests (mainly spruce-dominated production stands, combined with natural and semi-natural forest remnants), agricultural habitats (dominated by managed meadows), and urban areas.

We studied plant–pollinator interactions in the understorey of four semi-natural forest sites situated along the elevational gradient from the foothills to near the natural timberline (450, 600, 800, and 1,000 m a.s.l.; Table 1; Fig. 1). Using specialised forest maps provided by the conservation authorities, we selected forest stands with a semi-natural tree species composition characteristic of the respective elevation and region (Table 1). At each elevation, we established six non-overlapping transects (10 × 200 m). The transects were positioned to capture local vegetation heterogeneity, with deliberate inclusion of flower-rich understorey patches to ensure adequate recording of plant– pollinator interactions during the sampling period. Sampling was conducted during the early-spring phenological peak, in late April 2019 at the two lower elevations and early May 2020 at the two higher elevations, before full development of the deciduous canopy and at peak understorey flowering (Kudo et al. 2008) and forest pollinator activity (Harrison et al. 2018).

**Figure 1.**
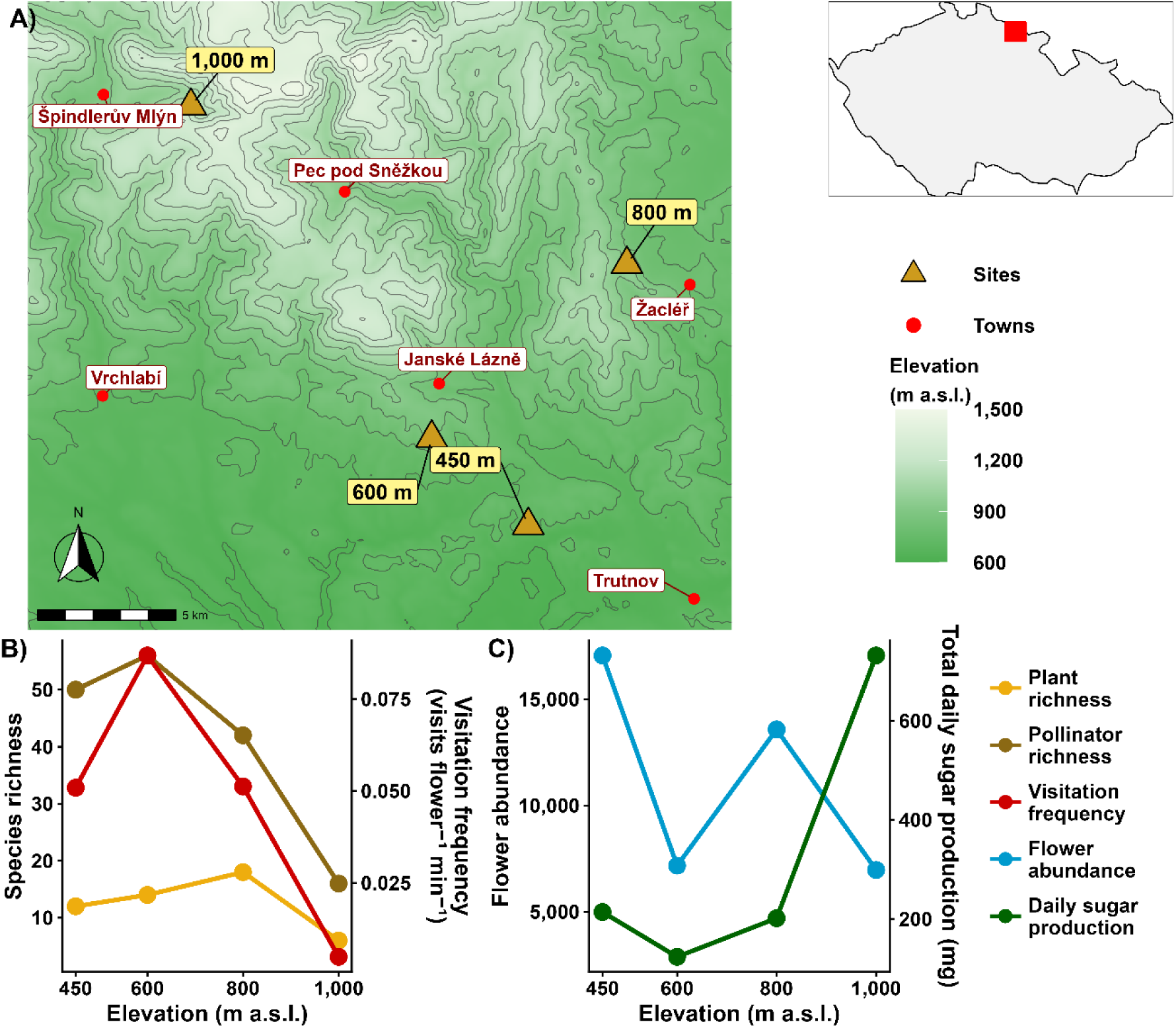
Study sites and broad elevational patterns in plant–pollinator communities in the Krkonoše Mountains, Czechia. (A) Locations of the four forest sites along the elevational gradient. (B) Plant and pollinator species richness and total visitation frequency of pollinators at each elevation. (C) Total flower abundance and total daily nectar sugar production at each elevation.

**Table 1.** Locations, habitat characteristics, species richness, visitation frequency and plant– pollinator network structure at four forest sites along the elevational gradient in the Krkonoše Mountains, Czechia.

| **Elevation (m a.s.l.)** | **450 m** | **600 m** | **800 m** | **1,000 m** |
| --- | --- | --- | --- | --- |
| **Site name** | Hradecek | Janske Lazne | Zacler | Spindleruv Mlyn |
| **Coordinates** | 50.5849°N,<br>15.8276°E | 50.6134°N,<br>15.7779°E | 50.6699°N,<br>15.8782°E | 50.7214°N,<br>15.6543°E |
| **Habitat** | Lowland mixed deciduous forest | Closed-canopy beech forest | Mixed beech-spruce forest | Montane sparse-canopy spruce forest |
| **Flowering plant species richness** | 12 | 14 | 18 | 6 |
| **Visited plant species richness** | 11 | 13 | 17 | 6 |
| **Pollinator species richness** | 50 | 56 | 42 | 16 |
| **Visitation frequency of all pollinators** (visits × flower<sup>-1</sup> × min<sup>-1</sup>) | 0.059 | 0.099 | 0.061 | 0.009 |
| **Visitation frequency of pollinator species** (visits × flower<sup>-1</sup> × min<sup>-1</sup>) | 0.051 | 0.087 | 0.051 | 0.005 |
| **Flower abundance** | 17,057 | 7,169 | 13,578 | 6,964 |
| **Total daily nectar sugar production (mg)** | 213.82 | 123.38 | 201.34 | 731.61 |
| **Network characteristics** |  |  |  |  |
| **Realised links** | 119 | 169 | 102 | 23 |
| **Connectance** | 0.216 | 0.232 | 0.143 | 0.240 |
| **Network-level specialisation $H_2'$** | 0.628 | 0.404 | 0.793 | 0.801 |
| **Weighted modularity $Q$** | 0.523 | 0.362 | 0.683 | 0.540 |
| **Nestedness $NODF$** | 34.71 | 38.49 | 22.56 | 15.09 |
| **Mean plant species-level specialisation $d'$ (±SD)** | 0.554 (±0.154) | 0.440 (±0.146) | 0.655 (±0.220) | 0.804 (±0.116) |
| **Mean pollinator species-level specialisation $d'$ (±SD)** | 0.377 (±0.148) | 0.337 (±0.178) | 0.547 (±0.177) | 0.475 (±0.253) |

### Flowering plants and floral traits

We first identified all plant species known or expected to receive insect visits that were flowering within the studied transects during the sampling period. Flower abundance was quantified once at each elevation, approximately midway through the corresponding pollinator recording period. We counted the flowers of each species across all six transects directly whenever feasible. For species with flower-rich inflorescences, we counted all inflorescences and estimated the mean number of flowers per inflorescence from at least 30 representative inflorescences, and their total flower abundance was then calculated by multiplying these two numbers.

We measured flower size and floral-tube length using digital callipers. Whenever possible, at least 15 flowers from at least five individuals were measured per species. For rare species, we measured at least five flowers, each from a different individual. The measurements were pooled across all elevations at which a species occurred, providing one trait value per species without considering intraspecific variation among elevations. Flower size was defined as the maximum linear dimension of the floral projection in frontal view. Floral-tube length was defined functionally as the distance that a pollinator had to traverse to access the nectar. Depending on flower morphology, this included the corolla tube, floral spurs, and any adjacent sepals contributing to the functional tube.

Daily nectar sugar production was estimated from at least 30 apparently fresh flowers per species, sampled across all elevations, following Janeček et al. (2021). The flowers were enclosed in appropriately sized mesh bags for 24 h to prevent nectar depletion by pollinators. The accumulated nectar was subsequently collected using microcapillary tubes or extracted by washing with distilled water using Hamilton syringes, depending on flower size and morphology. Samples were stored in vials containing distilled water at −18 °C and subsequently dried at 50 °C. The total mass of nectar sugars was quantified using capillary zone electrophoresis with contactless conductivity detection following Vlčková et al. (2023). The sugar mass measured after 24 h was averaged across the analysed flowers of each species and used as an estimate of daily nectar sugar production per flower (mg flower⁻¹ day⁻¹).

For each elevation, community means of flower size, floral-tube length, and daily sugar production per flower were calculated from species-level means weighted by the estimated flower abundance of each species. Total daily nectar sugar production was estimated by multiplying the mean daily sugar production per flower by the flower abundance of each species and summing the resulting values across all species.

### Pollinator recording and identification

We video-recorded flower visitors to all selected plant species flowering within the transects during the study period. Whenever possible, seven individuals of each plant species at each elevation were monitored for 24 h using VIVOTEK IB8367-T security cameras equipped with infrared night vision (for details, see Mertens et al. 2021; Klomberg et al. 2022). When a plant species was too rare within the transects to provide sufficient focal individuals, additional individuals were recorded in their surroundings. The cameras were positioned approximately 0.5–1 m from the focal flowers and camouflaged. Recording was suspended on days when heavy rainfall was forecast, whereas recordings containing shorter periods of light rain were retained because such conditions are typical of early spring in the temperate zone.

Recorded flower visitors were detected either using semi-automated motion detection with MotionMeerkat 2.0.5 (Weinstein 2015), when recording conditions allowed, or through manual review at accelerated playback speed. Only visitors that clearly touched the anthers and/or stigmas of the recorded flowers were considered potential pollinators and included in the analyses; hereafter, we refer to them as *pollinators*. A visit was defined as a single continuous presence of a pollinator in the recorded frame during which it contacted the anthers and/or stigmas of at least one flower; disappearances from the frame lasting ≤2 s were considered part of the same visit. For each visit, we recorded the number of individual flowers whose reproductive organs were contacted.

Pollinators were assigned to nine functional groups: bumblebees, honeybees, other bees, other hymenopterans, hoverflies, other flies, beetles, moths and butterflies. Records outside these groups were excluded from the analyses. Within the nine groups, pollinators were identified by experienced entomologists to the best possible taxonomic resolution, often as morphospecies (a morphologically distinct group of pollinators to which individuals could be assigned consistently across recordings, plant species and elevations, with reasonable confidence that they represented the same biological species, although formal species-level identification was not possible). Hereafter, the term *pollinator species* includes both species and morphospecies.

### Data processing and network analysis

All data preparation and analyses were performed in R v. 4.6.1 (R Core Team 2026). To account for differences in the number of recorded flowers and recording duration, sampling effort was quantified separately for each plant species at each elevation as flower-minutes. This measure represents the summed time for which individual flowers of each plant were recorded at each elevation. Visitation frequency was then calculated as the total number of flowers visited divided by the corresponding sampling effort and expressed as flowers visited per recorded flower per minute. Total visitation frequency at each elevation was calculated by summing visitation frequencies across the corresponding plant–pollinator interactions.

Elevational turnover of pollinator functional groups’ proportions (Fig. 2, Table S2) included all pollinators assigned to one of the nine functional groups, irrespective of the taxonomic resolution of their identification. All other summaries, figures, and analyses were restricted to pollinator species.

**Figure 2.**
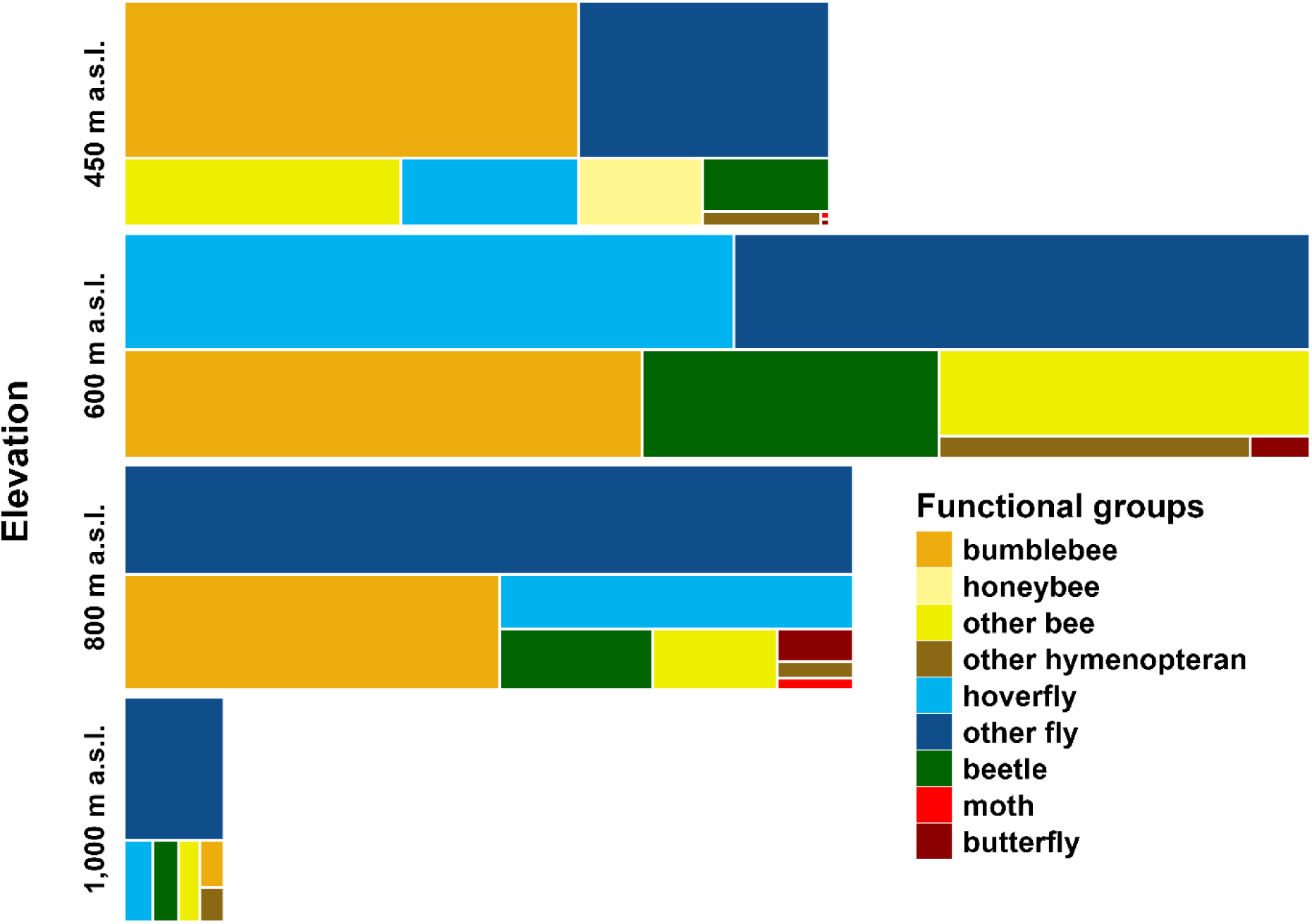
Composition and total visitation frequency of pollinator functional groups at four elevations in the Krkonoše Mountains, Czechia. The figure includes all pollinators from the nine functional groups that contacted the anthers and/or stigmas of the recorded flowers, irrespective of the resolution of their taxonomic identification. Within each elevation, rectangle area is proportional to the summed visitation frequency of the corresponding functional group. The total width occupied by the rectangles is proportional to total visitation frequency at that elevation. Visitation frequency is expressed as visits × flower⁻¹ × min⁻¹. Numerical values and percentages are provided in Table S2.

We constructed a separate quantitative plant–pollinator interaction matrix for each elevation, with each cell containing the summed visitation frequency of the corresponding plant–pollinator interaction. Only species involved in at least one retained interaction were included. Consequently, the number of plant species in a network could be lower than the total species richness of flowering plants recorded in the transects. Before calculating quantitative specialisation indices and modularity, visitation frequencies were multiplied by 10⁸ and rounded down to integers. This transformation retained the relative magnitudes of the interaction frequencies and all realised links while reflecting the realised sampling effort and satisfying the integer-input requirements of the analytical functions.

We characterised each network by connectance, network-level specialisation (*H₂′*), weighted modularity (*Q*), nestedness (*NODF*), and mean species-level specialisation (*d′*) of plants and pollinators, using the bipartite package (Dormann et al. 2008). Connectance is the proportion of realised links among all possible plant–pollinator combinations. *H₂′* measures overall specialisation as the deviation of observed interaction frequencies from random partner choice and ranges from 0 (complete generalisation) to 1 (maximum specialisation). It accounts for differences in network size and species’ total interaction frequencies (Blüthgen et al. 2006). Weighted modularity describes the extent to which interaction frequency is concentrated within groups of plants and pollinators rather than distributed between them. It was calculated using the stochastic Dormann–Strauss algorithm, which searches for the division of the quantitative network that maximises modularity (Dormann and Strauss 2014). The search was repeated ten times for each network, and the highest *Q* was retained. *NODF* measures the extent to which the partners of less-connected taxa form subsets of those of more-connected taxa and was calculated from the binary interaction structure (Almeida-Neto et al. 2008). Species-level *d′*, analogously to *H₂′*, ranges from 0 (generalists) to 1 (specialists) (Blüthgen et al. 2006). For each network, we calculated the arithmetic mean and SD of *d′* across all plant taxa and, separately, across all pollinator taxa.

Elevational differences in network-level indices were evaluated descriptively because each elevation was represented by a single study site, with interpretation focused primarily on broad contrasts between elevations. Species-level *d′* values were compared among elevations separately for plants and pollinators using Kruskal–Wallis tests. Significant overall differences were followed by pairwise Dunn tests implemented directly in R, with Benjamini–Hochberg adjustment applied separately to the six plant and six pollinator comparisons.

## Results

Across the four elevations, we obtained 280 focal recordings of 26 flowering plant species, totalling 6,725.5 h (i.e. >9 months). The recordings contained 4,003 pollinator visits assigned to the nine functional groups. Most visits were made by other flies (1,770 visits, 44.2%) and bumblebees (879 visits, 22.0%), followed by hoverflies (497 visits, 12.4%) and beetles (403 visits, 10.1%). Of all recorded visits, 3,157 were assigned to 87 pollinator species and retained for the species-level analyses.

Species richness and visitation frequency peaked at mid-elevations, but the peak elevations differed. Flowering plant species richness was highest at 800 m, whereas pollinator species richness and visitation frequency were highest at 600 m. All three characteristics were lowest at 1,000 m, particularly visitation frequency, which was substantially lower than at the other elevations (Fig. 1B; Table 1). Flower abundance showed no consistent contrast between the two lower and two higher elevations. Despite low plant species richness and low flower abundance, total daily nectar sugar production was markedly higher at 1,000 m than at the remaining sites (Fig. 1C).

The estimated floral abundance was dominated by *Mercurialis perennis* at 450 m (12,232 flowers, 71.7% of all censused flowers), 600 m (2,742, 38.2%) and 800 m (7,997, 58.9%). *Chrysosplenium alternifolium* was also abundant at 600 m (2,175 flowers, 30.3%), as was *Corydalis cava* at 800 m (2,794, 20.6%). At the highest elevation, *Vaccinium myrtillus* accounted for 6,190 flowers, representing 88.9% of the local flower abundance.

Floral communities were characterised by different dominant species at each elevation. The largest numbers of flowers visited by pollinators were recorded for *Lathraea squamaria* at 450 m (982 flowers) and at 600 m (614), on *Adoxa moschatellina* at 800 m (807), and *C. alternifolium* at 1,000 m (131). After accounting for sampling effort, the highest visitation frequencies were observed for *L. squamaria* at 450 m (0.0139 visits **×** flower⁻¹ **×** min⁻¹), *Ficaria verna* at 600 m (0.0352), *A. moschatellina* at 800 m (0.0203), and *C. alternifolium* at 1,000 m (0.0036). No flowers of *Asarum europaeum* were visited at 600 or 800 m, although seven were visited at 450 m. Similarly, no flowers of *Viola reichenbachiana* were visited at 600 m, compared with three visited at 800 m.

Among pollinators retained in the species-level analyses, *Bombus pascuorum* (Hymenoptera) was the most frequently recorded species overall, with 462 visits, representing 14.6% of all retained visits. It was followed by Nematocera sp. “striped” morphospecies (Diptera; 422 visits, 13.4%) and *Bombus pratorum* (Hymenoptera; 221 visits, 7.0%). At 450 m, *B. pratorum* and *B. pascuorum* together contributed almost half of the total network visitation frequency. *B. pascuorum* was also the most important pollinator at 600 and 800 m, contributing 15.1% and 25.3% of the respective network visitation frequencies; at 800 m, the Nematocera sp. “striped” contributed a further 23.5%. The much smaller network at 1,000 m was not dominated by a single pollinator species, with *Bombyliidae* sp. 1 (Diptera) contributing the largest share of its visitation frequency (18.7%; Fig. 3).

**Figure 3.**
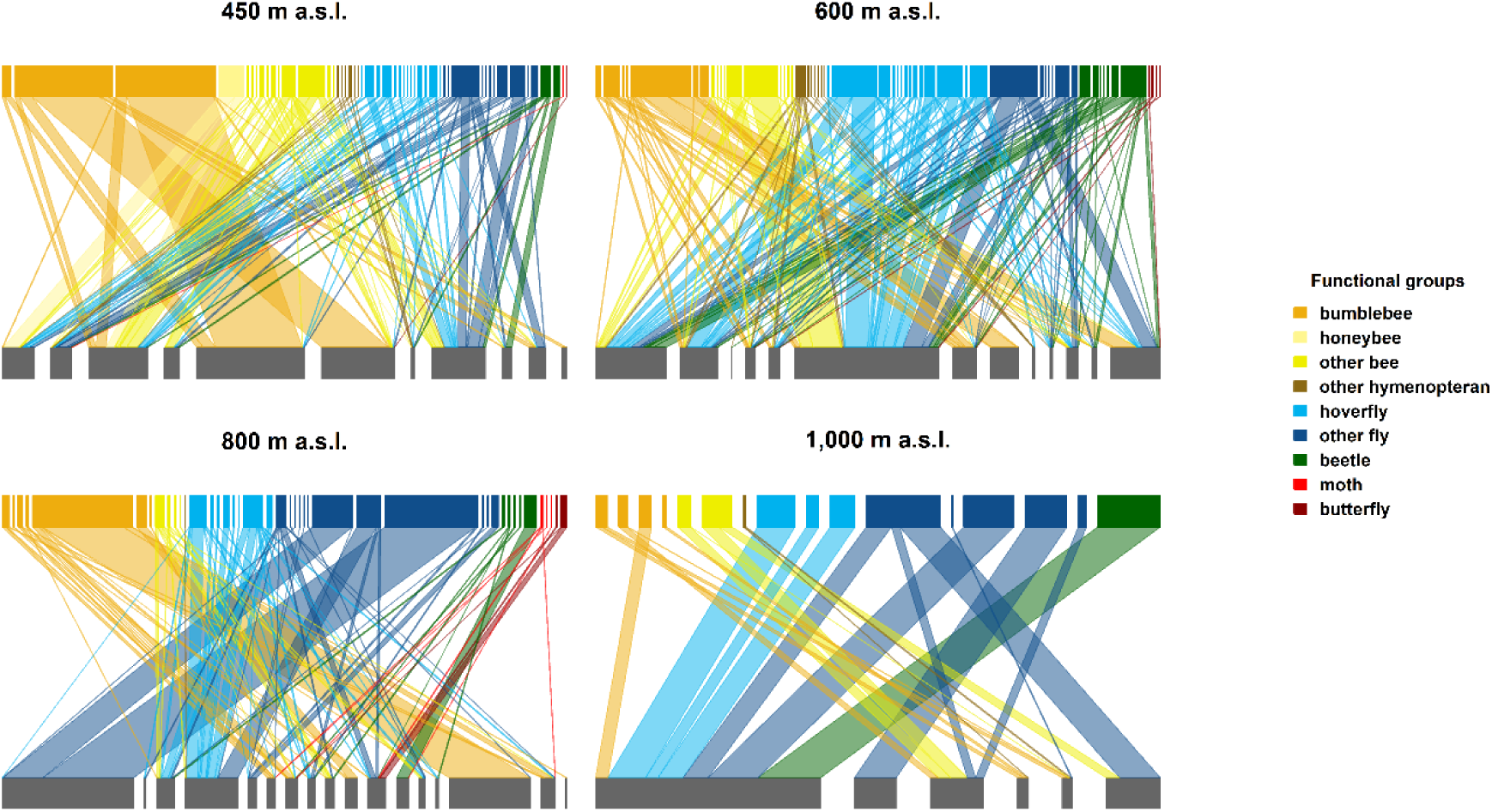
Quantitative plant–pollinator networks at four elevations in the Krkonoše Mountains, Czechia. Only visits involving contact with the floral reproductive organs and pollinators identified to species or morphospecies are included. Lower grey nodes represent plant species and upper nodes represent pollinator species, coloured according to pollinator functional group. Box widths and connecting-link widths are proportional to summed visitation frequencies.

### Elevational turnover in pollinator functional groups

The relative contributions of pollinator functional groups to total visitation frequency changed considerably among elevations (Fig. 2; Table S2). Hymenopterans accounted for 35.6–63.3% of total visitation frequency at the two lower elevations, compared with 16.6–31.8% at the two higher elevations. Conversely, the contribution of dipterans increased from 32.4–51.8% at the lower elevations to 60.5–74.2% at the higher elevations. Bumblebees had the highest visitation frequency at 450 m, whereas visitation frequency at 600 m was distributed mainly among hoverflies, other flies and bumblebees. Other flies had the highest visitation frequency at both 800 m and 1,000 m.

Visitation frequencies of the individual bee groups showed contrasting elevational patterns. Bumblebee visitation frequency decreased progressively with elevation, visitation by other bees peaked at 600 m, and honeybee visitation was recorded only at 450 m. Beetle visitation frequency likewise peaked at 600 m and was lowest at 1,000 m. Lepidopteran visitation frequencies remained low at all elevations, together accounting for no more than 2.1% of total visitation frequency (Fig. 2; Table S2).

### Network composition and structure

The plant–pollinator network at 1,000 m was substantially smaller than the other three networks and contained only 23 realised links, compared with 102–169 links at the other elevations (Fig. 3; Table 1). The two higher-elevation networks had higher network-level specialisation (*H₂′*) and lower nestedness (*NODF*) than the two lower-elevation networks. Weighted modularity (*Q*) was highest at 800 m and lowest at 600 m, whereas the values at 450 m and 1,000 m were similar. Connectance showed no clear contrast between the lower and higher elevations (Fig. 4A; Table 1).

**Figure 4.**
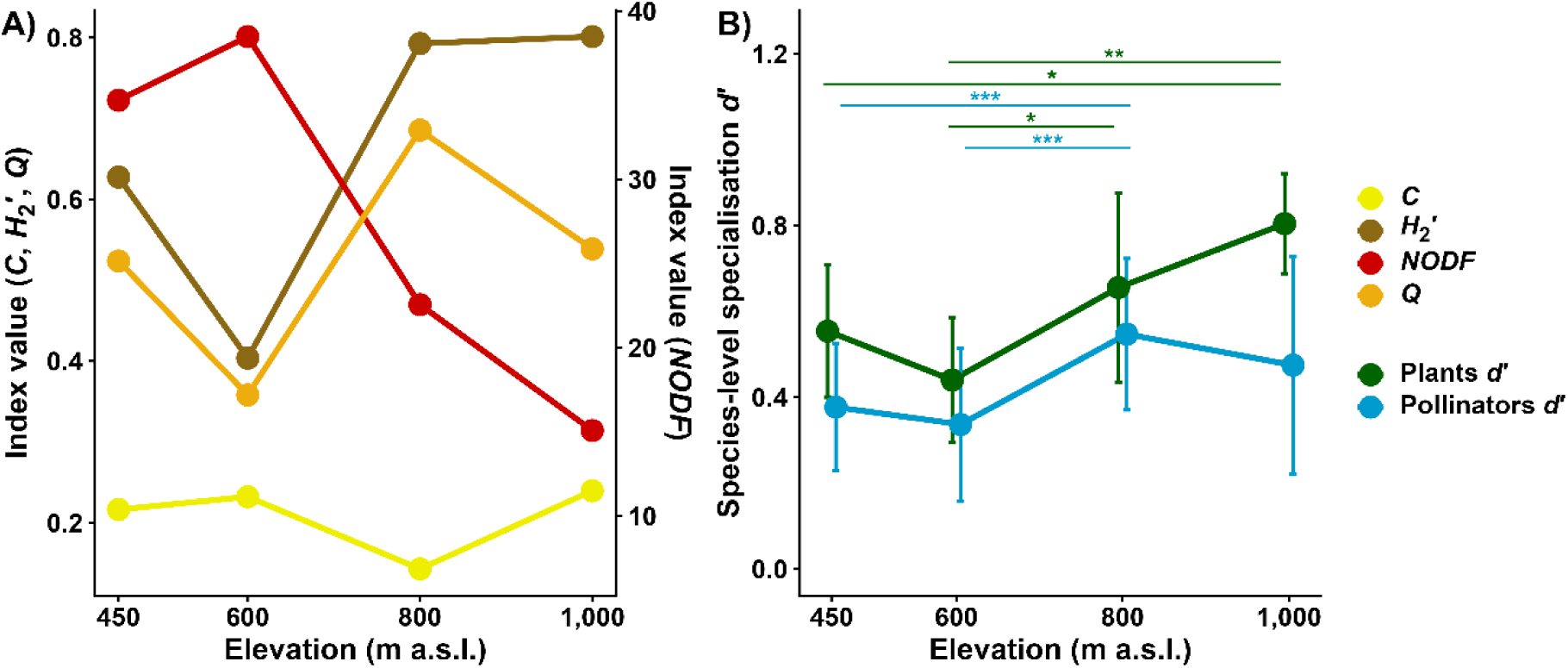
Plant–pollinator network structure and species-level specialisation along the elevational gradient in the Krkonoše Mountains, Czechia. (A) Connectance (C), network-level specialisation (H₂′), weighted modularity (Q), and binary nestedness (NODF) of the four pollination networks. (B) Mean species-level specialisation (d′) of plants and pollinators; error bars indicate ±SD among species within each network. Horizontal brackets show significant pairwise Dunn comparisons (∗ P<0.05,∗∗ P<0.01,∗∗∗ P<0.001) following significant Kruskal–Wallis tests, with Benjamini–Hochberg adjustment applied separately to plants and pollinators. Complete pairwise results are provided in Table S1.

Mean species-level specialisation (*d’*) differed among elevations for both plants (Kruskal– Wallis: H=16.05, df=3, P=0.001) and pollinators (H=32.47, df=3, P<0.001). Plant *d′* was significantly higher at 1,000 m than at both 450 and 600 m, and higher at 800 m than at 600 m. Pollinator *d′* was significantly higher at 800 m than at both 450 and 600 m. None of the remaining pairwise comparisons were significant after the Benjamini–Hochberg adjustment (Fig. 4B; Table S1).

### Floral traits

Community-weighted mean flower size varied among elevations without a consistent contrast between the two lower and two higher sites (Fig. 5A). In contrast, community-weighted mean floral tube length consistently increased with elevation (Fig. 5B). Community-weighted daily nectar sugar production per flower was similar at 450, 600 and 800 m but substantially higher at 1,000 m (Fig. 5C). Correspondingly, total daily sugar production reached 731.6 mg at 1,000 m, compared with 123.4–213.8 mg at the other elevations (Fig. 1C). These high values at the highest elevation were predominantly driven by the abundant flowers and relatively high nectar sugar production of *Vaccinium myrtillus*, which accounted for 98.9% of the estimated daily sugar production at 1,000 m.

**Figure 5.**
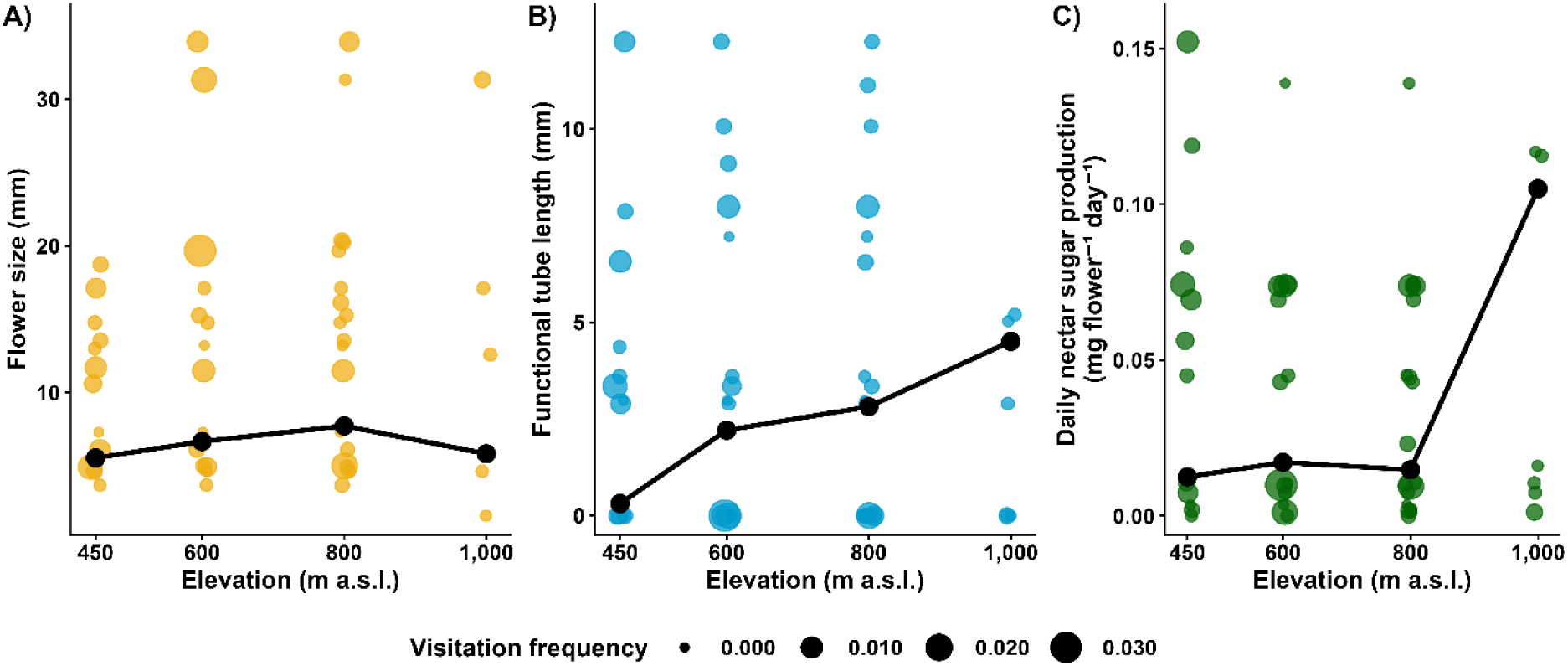
Community-weighted means of floral traits and daily nectar sugar production of plant species along the elevational gradient in the Krkonoše Mountains, Czechia. Coloured points represent plant species occurring at each elevation, with their size proportional to the visitation frequency. Black points and connecting lines represent community means weighted by flower abundance of individual species. (A) Flower size, (B) functional floral-tube length, and (C) daily nectar sugar production per flower.

## Discussion

Our results show that early-spring forest plant–pollinator communities changed along the elevational gradient through partly independent responses of plant resources, pollinator composition and interaction structure, rather than through a uniform change in all community properties. Partly confirming our expectations, plant and pollinator species richness and visitation frequency were lowest at the highest site, although their differing maxima at intermediate elevations demonstrated that plants and pollinators did not respond synchronously. Similar mismatches between the elevational patterns of pollinators and their floral hosts have been reported in other non-tropical mountains (Sponsler et al. 2022; Minachilis et al. 2023), while similar mid-elevation maxima in interacting species richness and interaction frequency were found by Adedoja et al. (2018). These results fit the prevailingly non-linear elevational patterns in species richness reviewed by Guo et al. (2013) and Dolson and Kharouba (2024). Flower abundance showed no consistent contrast between the lower and upper elevations, differing from the elevational decline reported by Aguirre and Junker (2024) but agreeing with the spatial and seasonal variation in flower abundance found by Lefebvre et al. (2018). The observed abundance patterns were strongly affected by dominant species, particularly *M. perennis* at the three lower sites and *V. myrtillus* at the highest site. The latter species was also responsible for nearly all estimated nectar sugar production at 1,000 m. Consequently, the unexpectedly high total sugar production at the highest elevation represented local dominance by a single plant rather than a general increase in community floral resources. Comparable community-level information on nectar sugar production remains largely unavailable from temperate elevational gradients.

Confirming our expectation, pollinator communities shifted from a greater contribution of hymenopterans at lower elevations towards increasing dominance by dipterans at the upper sites (Fig. 2; Table S2). This agrees with patterns reported from other non-tropical mountain systems (Kearns 1992; Lefebvre et al. 2018; McCabe et al. 2019; McCabe and Cobb 2021). Nevertheless, individual functional bee groups did not show this general elevational response, corresponding with the contrasting inter-group patterns reported elsewhere (Baumann et al. 2021; Sponsler et al. 2022; Sommaggio et al. 2022; Minachilis et al. 2023). Most importantly, the increasing contribution of dipterans in our study was driven predominantly by other flies rather than hoverflies, which can be an important part of pollination networks but remain comparatively neglected (Orford et al. 2015). Their prominence in our recordings suggests that they may be important components of early-spring forest pollination, although we did not measure pollination effectiveness.

The lack of elevational differences in flower size agrees with community-level studies from the Swiss and Austrian Alps (Fabbro and Körner 2004; Junker and Larue-Kontić 2018). In contrast, the increasing floral tube length towards the upper sites differs from the absence of such a pattern in the Austrian Alps (Junker and Larue-Kontić 2018), further demonstrating that elevational changes in floral characteristics are trait- and community-dependent. The increasing representation of plants with longer floral tubes could alter the accessibility of floral resources because morphological barriers can restrict flower use by particular visitors (Santamaría and Rodríguez-Gironés 2007). However, without corresponding pollinator traits, we cannot determine whether this compositional change contributed to the observed differences in interactions or specialisation.

We consider the increase in specialisation towards the upper sites as our most interesting result because it contrasts with both the altitudinal niche-breadth hypothesis and most comparable elevational studies from non-tropical mountains. Contrary to our expectation, the upper-elevation communities were more specialised at both network and species levels, whereas decreasing specialisation or increasing generalisation has been reported from alpine grasslands and the only previously studied temperate forest gradient (Hoiss et al. 2015; Chesshire et al. 2021; Lara-Romero et al. 2019). Together with the non-linear responses found by Aguirre and Junker (2024), these contrasting outcomes support the broader conclusion that climatic patterns in plant–pollinator specialisation are strongly dependent context-dependent (Sakhalkar et al. in press). The coincidence of low plant species richness and high specialisation at the highest elevation is consistent with the association between lower plant diversity and stronger network specialisation reported by Schleuning et al. (2012). The high specialisation at 1,000 m partly reflected its uneven and strongly partitioned network: *A. nemorosa* accounted for 56.9% of total network visitation frequency, with six of its seven pollinator species recorded from no other plant, while *C. alternifolium* and Nematocera sp. “striped” formed an isolated pair. Possible, non-exclusive mechanisms for such specialised interactions include uneven resource and species-abundance distributions, morphological restrictions on flower use, tighter temporal matching within shorter flowering and foraging periods, and physiological filtering of pollinator groups with different climatic sensitivities (Santamaría and Rodríguez-Gironés 2007; Kaiser-Bunbury et al. 2014; Glaum et al. 2021; Sakhalkar et al. in press). Our data cannot distinguish among these mechanisms.

The other network characteristics showed less consistent elevational patterns. Lower nestedness at the upper sites contrasted with the increase reported by Chesshire et al. (2021), while modularity showed no clear elevational pattern. Such variation agrees with the context-dependent network responses reported by Aguirre and Junker (2024) and may partly reflect geometric constraints on species co-occurrence along environmental gradients (Fibich et al. 2026). Because modularity is also influenced by network dimensions (Olesen et al. 2007), and each elevation was represented by a single network, these characteristics are best considered descriptive properties of the four local communities rather than evidence of general elevational patterns.

By providing detailed community-wide data from a temperate forest elevational gradient, this study fills an important geographic and habitat gap in plant–pollinator research. Each elevation was represented by six transects, but their aggregation into one network per site necessarily limits generalisation beyond the studied gradient. Nevertheless, descriptive data from poorly represented ecosystems are important both for documenting regional natural history and for broader comparative analyses, as demonstrated by the inclusion of these networks in recent global and elevational syntheses (Sakhalkar et al. in press; Fibich et al. 2026). The upper-elevational communities combined lower pollinator activity and greater dominance by non-syrphid flies with stronger quantitative partitioning of interactions. Their unusual increase in specialisation deserves particular attention, although replicated studies across other non-tropical forest gradients will be needed to determine whether it represents a more general elevational pattern in forest ecosystems or results from other characteristics of the studied communities. More broadly, our findings illustrate the problems of generalising elevational patterns from the limited range of open habitats and ecosystems represented in most existing studies.

## Acknowledgements

We are grateful to Lucas Brisson, Fotoula Papandreou, Daniel Souto Vilarós, and Guillermo Uceda-Gómez for their help with fieldwork, to Ivan Šonský and Fabian Simon Klimm for help with processing the video recordings, and to the staff of the Krkonoše National Park, mainly Stanislav Březina and Robin Böhnish, for their help with site selection and various logistical matters. Our research in the Krkonoše NP was performed under the required permits. This study was funded by the Czech Science Foundation (21-24186M).

## Author contribution

RT and ŠJ conceived and designed the study; RT supervised the study; EC, AS, SPS, and INK led the fieldwork and data processing; AS, ŠJ, SPS, INK, EC, DA, SD, KJ, and YK sampled the data and material; AS, SPS, INK, KJ, YK, and JEJM processed the video recordings; JH, JF, SPS, and RT identified flower visitors; TK, RP, and KJ analysed nectar composition; RT, AS, SPS, and DA processed and analysed the data and prepared visualisations; RT wrote the first draft based on remarks from AS; all authors edited the manuscript and approved its submission.

## Conflict of interest

The authors declare no conflict of interest.

## AI disclosure statement

ChatGPT (GPT-5.5 and GPT-5.6 Sol, OpenAI) was used to improve English expression and assist with checking R scripts used for data preparation and analysis. All suggestions and outputs were critically reviewed and verified by the corresponding author, who takes full responsibility for these.

---

**Table S1.** Pairwise comparisons of species-level specialisation (d′) of plants and pollinators along the elevational gradient in the Krkonoše Mountains, Czechia. Pairwise differences were evaluated using Dunn tests following significant Kruskal–Wallis tests. *P* values were adjusted using the Benjamini– Hochberg procedure separately for plants and pollinators. Significant adjusted *P* values are shown in bold, with asterisks indicating significance levels (*P < 0.05, **P < 0.01, ***P < 0.001).

| Compared elevations (m a.s.l.) | Plants |  | Pollinators |  |
| --- | --- | --- | --- | --- |
| | Z | adjusted $P$-values | Z | adjusted $P$-values |
| 450 × 600 | 1.265 | 0.206 | 1.164 | 0.244 |
| 450 × 800 | −1.302 | 0.206 | −4.227 | ***<0.001 |
| 450 × 1,000 | −2.573 | *0.020 | −1.378 | 0.202 |
| 600 × 800 | −2.774 | *0.017 | −5.445 | ***<0.001 |
| 600 × 1,000 | −3.695 | **0.001 | −2.195 | 0.056 |
| 800 × 1,000 | −1.689 | 0.137 | 1.664 | 0.144 |

**Table S2.** Visitation frequencies of pollinator functional groups along the elevational gradient in the Krkonoše Mountains, Czechia. Each cell gives the percentage of the total visitation frequency followed in parentheses by the summed visitation frequency (visits × flower⁻¹ × min⁻¹) at the respective elevation. Values include all recorded pollinators assigned to the nine functional groups, irrespective of taxonomic resolution of pollinator identification. The four pollinator order values presented at the table bottom are derived summaries of the corresponding functional groups.

|  | 450 m | 600 m | 800 m | 1,000 m |
| --- | --- | --- | --- | --- |
| **Pollinator functional group** |  |  |  |  |
| **Bumblebees** | 45.00%<br>(0.02654) | 21.03%<br>(0.02077) | 26.36%<br>(0.01608) | 5.18%<br>(0.00046) |
| **Honeybees** | 5.30%<br>(0.00314) | 0.0%<br>(0) | 0.0%<br>(0) | 0.0%<br>(0) |
| **Other bees** | 11.86%<br>(0.00700) | 12.03%<br>(0.01189) | 4.62%<br>(0.00282) | 7.67%<br>(0.00068) |
| **Other hymenopterans** | 1.09%<br>(0.00064) | 2.54%<br>(0.00251) | 0.79%<br>(0.00048) | 3.72%<br>(0.00033) |
| **Hoverflies** | 7.60%<br>(0.00448) | 26.66%<br>(0.02633) | 11.70%<br>(0.00713) | 10.40%<br>(0.00092) |
| **Other flies** | 24.78%<br>(0.01461) | 25.18%<br>(0.02487) | 48.78%<br>(0.02975) | 63.83%<br>(0.00568) |
| **Beetles** | 4.26%<br>(0.00251) | 12.06%<br>(0.01191) | 5.70%<br>(0.00348) | 9.20%<br>(0.00082) |
| **Moths** | 0.04%<br>(0.00003) | 0.0%<br>(0) | 0.53%<br>(0.00032) | 0.0%<br>(0) |
| **Butterflies** | 0.04%<br>(0.00002) | 0.49%<br>(0.00048) | 1.52%<br>(0.00093) | 0.0%<br>(0) |
| **Total visitation frequency** | **0.05898** | **0.09876** | **0.06099** | **0.00889** |
| **Pollinator order** |  |  |  |  |
| **Hymenoptera** | 63.28%<br>(0.03732) | 35.60%<br>(0.03516) | 31.77%<br>(0.01938) | 16.57%<br>(0.00147) |
| **Diptera** | 32.38%<br>(0.01910) | 51.84%<br>(0.05120) | 60.47%<br>(0.03688) | 74.23%<br>(0.00660) |
| **Coleoptera** | 4.26%<br>(0.00251) | 12.06%<br>(0.01191) | 5.70%<br>(0.00348) | 9.20%<br>(0.00082) |
| **Lepidoptera** | 0.08%<br>(0.00005) | 0.49%<br>(0.00048) | 2.05%<br>(0.00125) | 0.0%<br>(0) |

## Notes

### Competing Interest Statement

The authors have declared no competing interest.

